# A lone spike in blood glucose can enhance the thrombo-inflammatory response in cortical vessels

**DOI:** 10.1101/2022.08.10.503527

**Authors:** Iftach Shaked, Conrad Foo, Rui Liu, Yingying Cui, Xiang Ji, Thomas Broggini, Philipp Mächler, Prithu Sundd, Anna Devor, Beth Friedman, David Kleinfeld

## Abstract

How transient hyperglycemia contributes to cerebro-vascular disease has been a challenge to study under controlled conditions. We present an approach to model luminal vessel thrombo-inflammation using amplified, ultrashort laser-pulses to physically disrupt brain-venule endothelium. Vessel disruption in conjunction with transient hyperglycemia from a single injection of metabolically active *D*-glucose results in real-time responses to venule damage that include rapid serum extravasation, platelet aggregation, and neutrophil recruitment, in normal mice. In contrast, vessel thrombo-inflammation following laser-induced vessel disruption is significantly reduced in mice injected with metabolically inert L-glucose. Thrombo-inflammation is pharmacologically ameliorated by a platelet inhibitor, by a scavenger of reactive oxygen species, or by a nitric oxide donor. For comparison, in diabetic mice injured vessel thrombo-inflammatory responses are also reduced by restoration of normo-glycemia. Our approach provides a controlled method to probe synergies of transient metabolic and physical vascular perturbations and reveals new aspects of brain pathophysiology.

## INTRODUCTION

Transient hyperglycemia secondary to metabolic demands and various stressors has been linked to adverse ischemic stroke outcomes (Fuentes et al., 2020; Gonzalez-Moreno et al., 2014; Yoon et al., 2017) and long-lasting effects on the patency of vessels. (El-Osta et al., 2008). While diabetes, or chronic hyperglycemia, are well defined in both human patients and mice models, transient hyperglycemia is only more recently appreciated and studied (Hall et al., 2018). Continuous glucose monitoring in humans, in comparison to single time-point monitoring, reveals that healthy individuals display high glucose transient spikes that reach prediabetic and even diabetic levels. These spikes in blood glucose concentration may occur either from rapid, high caloric intake or from an endocrine imbalance that can transiently be induced by different stressors including inflammation (Qing et al., 2020) and chronic social stress (van der Kooij et al., 2018). As a matter of public health, spikes in blood glucose concentration were suggested to play a role in activation of oxidative stress compromising vascular health (Monnier et al., 2006).

Is hyperglycemia, in the absence of comorbidities such as those present in people with diabetes, prothrombotic? Numerous results in model systems suggest a link between high glucose and a prothrombotic phenotype (Beckman et al., 2002). The formation of platelet-rich thrombi is initiated, at least in part, by soluble vascular stress signals such as thrombin, epinephrine, and adenosine diphosphate (Brzoska et al., 2020), as well as by exposure to collagen and laminin components of the extracellular matrix of a disrupted vessel (van der Meijden and Heemskerk, 2019). It is suggested that hyperglycemia increases platelet responsiveness to such agonists (Kraakman et al., 2001). Mechanistically, hyperglycemia increases production of mitochondrial reactive oxygen species (ROS) in primary endothelial cells in vitro (Nishikawa et al., 2000) and the constellation of ROS mediated pathogenic pathways provides a basis for the endothelial dysfunction observed in people with diabetes (Cai and Harrison, 2000; Geraldes et al., 2009; Ungvari et al., 2017). Importantly, blood fluidity is impacted by ROS through associated depletion of the vasoactive gaseous transmitter nitric oxide. This gas molecule is an essential cofactor for vasodilation (Lindauer et al., 1999) and further functions as a checkpoint for inhibition of platelet activation (Naseem and Roberts, 2011). Platelet activation in turn initiates an inflammatory cascade that is mediated, in part, by neutrophil recruitment (Sreeramkumar et al., 2014).

As a means to establish a potential link between a spike in blood glucose concentration in the blood stream and a prothrombotic phenotype, we introduce a new model of vascular tissue disruption. Our model makes use of targeted vessel disruption from amplified ultra-short laser pulses as a surrogate for intra or juxta-luminal vessel injuries that lead to platelet activation. In their pioneering work, the Furies used laser induced vascular disruption as a means to measure thrombus formation, in real time, within the cremaster artery of anesthetized rats (Falati et al., 2002). This mimics the pathophysiology of spontaneous vessel wall tissue disruption that may be variously triggered by high wall-shear stress (Versteeg et al., 2013), infection (Stark and Massberg, 2021), and aging (Le Blanc and Lordkipanidzé, 2019). Our experimental platform extends the Furie model to examine effects of transient hyperglycemia on tissue disruption in brain vasculature. Our approach features the use of focused optical excitation for vessel disruption to the unique environment of brain vasculature. This allows a controlled vessel disruption through the nonlinear activation of light and tissue and is free of thermal damage to the vessel wall (Loesel et al., 1998; Tsai et al., 2009) and so we can precisely induce single vessel tissue disruption with minimal collateral damage. We make use of a transcranial imaging so that brain homeostasis is unaffected (Drew et al., 2010), as well as the potential for inflammation induced by a craniotomy (Lagraoui et al., 2012). Importantly, these advances allow us to perform experiments on awake animals and thus avoid confounds from anesthesia. In particular, different anesthesia agents, i.e., fentanyl/fluanisone/midazolam, ketamine/xylazine, isoflurane, and pentobarbital, significantly influence glucose metabolism (Windeløv et al., 2016). The present study also extends the past use of such laser pulses to ablate individual blood vessels (Nishimura et al., 2010; Nishimura et al., 2006) and somata (Cheng et al., 2021; Koyama et al., 2016) in vivo.

We chose to target cortical cerebral veins for disruption based on four motivating factors. The first two are biomedical: (i) the central sinus and venous branches act as an immune hub, with both innate and adaptive immune activity (Rustenhoven et al., 2021); and (ii) cerebral venous thrombosis is an important cause for stroke especially in young adults (Silvis et al., 2017). The other two are technical: (iii) the anatomy and functions of the vein makes platelet activation and their interaction with neutrophils much more likely to occur in veins than in arteries (Nemeth et al., 2020; van der Meijden and Heemskerk, 2019); and (iv), pial veins are claimed to be topologically organized as a tree (Coelho-Santos and Shih, 2020), which facilitates independence among intravascular thrombi generated in different branches that drain to the sagittal sinus. This organization allows us to perform calibrated, repeated injuries by targeting independent venous segments while controlling blood glucose levels.

## RESULTS

To test whether hyperglycemia in isolation exacerbates a thrombo-inflammatory response to vascular tissue disruption, we examined responses in cortical veins of wild type mice where paired injuries are performed in a blinded fashion in the same subject under hyperglycemic (*D*-glucose) or normoglycemic (*L*-glucose) challenge. Pharmacological manipulations to blunt hyperglycemic pathogenic pathways in transiently hyperglycemic mice were tested for their effects on mitigation of the laser-triggered vascular tissue disruption. As controls, we utilized two models of murine diabetes to establish a baseline standard for the response to vascular tissue disruption in the presence of comorbidities linked to diabetes. All data was taken from pial venules above parietal cortex in adult mice.

### Cerebrovascular mapping

We first characterized our test bed by mapping the topology of the pial venule network. We filled the vasculature of mice with a fluorescent label and obtained overlapping fluorescent images of the superior sagittal sinus and the complete extent of venules that drain somatosensory cortex into the sinus (11 vessels across 5 mice; **Figure 1A**). In agreement with past claims, we find that all venules appear as trees; this is unlike the pial arterioles, which form loops (Blinder et al., 2010). There is a broad distribution of branching orders, with a mean at a branching order of 11 (**Figure 1A**). Going forward, we choose 3^rd^ or 4^th^ order branches as target venules. The role of these venules in waste drainage (Coelho-Santos and Shih, 2020) coupled with their heightened immune activity (Rustenhoven et al., 2021) makes them the brain’s prothrombotic Achilles heel.

**Figure 1.**
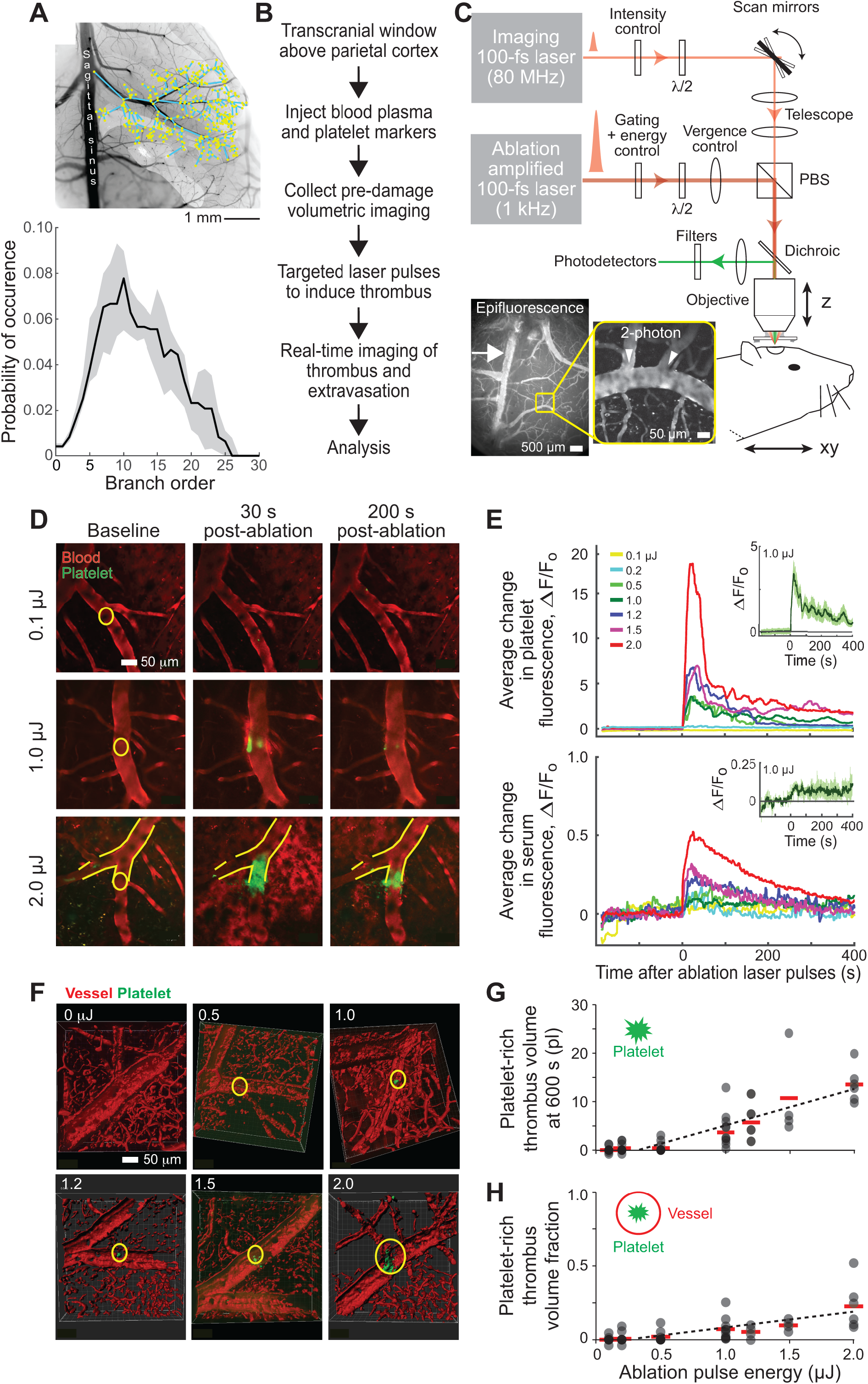
Sample preparation and calibration studies for cerebrovascular disruption by the targeted delivery of amplified ultra-short laser pulses. **(A)** Topology of the pial venules. The top diagram is taken from the reconstruction of vasculature in a single mouse. The bottom compilation incorporates 11 surface venules in five mice. **(B)** Time-line of study design. **(C)** Schematic representation of the optical system that combines two-photon laser scanning microscopy with optical ablation using a gated train of amplified ultra-short pulses. The inset is the image of surface vasculature of mouse cortex; arrow indicates branch of superior cerebral vein targeted for laser-induced disruption. **(D)** Fluorescent images of platelet (anti-CD41; green) and serum (Texas red dextran) fluorescence in cortical surface veins, before and 30 s and 200 s post irradiation using 0.1 µJ,1.0 µJ, and 2.0 µJ laser pulses. Focal region of the excitation spot is highlighted by the yellow circle. **(E)** Effects of varying laser energy, by increasing the power and keeping the number of pulses constant, on the time course of laser-induced aggregation of platelets and blood extravasation following the example data in panel D. Each curve is generated from the average change in vein fluorescence relative to baseline values (6 veins at one vein per animal at 0.1 µJ, 0.2 µJ, and 1.0 µJ; 3 for 1.2 µJ, and 1.5 µJ; 11 veins for 0.5 µJ; and 5 veins for 2.0 µJ); 40 mice total. Inset highlights the mean and standard error for the samples at 1.0 µJ; data was smoothed with 2.4 s median filter. **(F)** Representative three-dimensional reconstructions of platelets aggregates (green) within vasculature (red), collected 400 to 600 s after laser-induced vascular disruption. **(G)** Platelet-rich thrombi volumes for data collected between 400 and 870 s after laser-induced vascular disruption; we mark the midpoint time of ∼ 600 s post laser damage. Same veins as in panel E. The red bar is the mean. The responses at 0.5 µJ is not statistically different from zero (p = 0.10) while that at 1.0 µJ is different (p < 0.01). The data is fit with a threshold-linear function with intercept of 0.24 µJ and slope of 9.8 ± 2.4 pL/µJ. **(H)** Thrombus volume fractions, i.e., volume normalized to the volume of a sphere whose radius equals that of the vein; reanalysis of the data in panel G. The response at 0.5 µJ is not statistically different from zero (p = 0.18), while that at 1.0 µJ is different (p < 0.016). The data is fit with a threshold-linear function, with intercept forced to be 0.24 µJ, for a slope of 0.18 ± 0.03 µJ^-1^.

### Venule laser targeting

A challenge for in vivo perturbation of cerebral vasculature function is the design of a quantitatively reproducible perturbation. This requires a defined protocol (**Figure 1B**) and test metrics for the perturbation and the response. Vessel fluorescence was imaged from retro-orbital injection of Texas red dextran, while platelets were labeled by injection of AF594-conjugated anti-CD41 antibodies (**Figure 1C**). To visualize neutrophil/leukocyte responses to laser-induced disruption, we utilized transgenic LysM-eGFP mice where blood leukocytes are endogenously labeled through expression of eGFP.

Targeted laser-induced disruption to the lumen of a venule was delivered by amplified ultra-short laser pulses of known energy (Nishimura et al., 2006). Real time changes in fluorescence were continuously recorded and normalized to baseline pre-injury levels to quantify the dynamic effects of laser-induced disruption across a region of interest (ROI) centered around the blood vessel in a single imaging plane. We tested the effect of laser pulse energies ranging from 0.1 µJ to 2.0 µJ (**Figure 1D-H**). Laser-induced disruption led to a sharp peak in platelet aggregation, as inferred from the fluorescence of labeled platelets. The amplitude in fluorescence decayed from its peak value, post-disruption, towards a plateau value after 400 s of recovery (**Figure 1E**). We reconstructed the volume of the platelet-rich thrombi during the plateau period from successive optical sections, i.e., a “z-stack” (**Figure 1F,G**). Thrombus volumes increased with increasing of laser pulse energies. The first statistically significant change from zero volume occurs at an energy of 1.0 µJ (p < 0.01; **Figure 1G**), as noted above, for which we observe thrombi of 3.6 ± 1.2 pl (mean ± SE). As a normalized metric, we estimated the measured thrombus volume relative to that of the volume for a hypothetical sphere whose diameter equals the diameter of the vessel (**Figure 1H**). We designate this quantity as the “thrombus volume fraction”. This normalized volume is small, i.e., 0.072 ± 0.024, so that the thrombi we form are modest in size and well controlled. Lastly, we inferred the extent of serum extravasation from the fluorescence of extravascular labeled serum within seconds after irradiation (**Figure 1D,E**).

In total, these test data highlight the quantitative relation between laser pulse energies, as a model of the pathophysiology vessel wall disintegration (Hechler et al., 2009), and intra- and juxta-luminal responses to vascular tissue disruption. It sets the stage for the use of this targeted assay to assess the increased susceptibility of vessels to irradiation damage by hyperglycemia.

### Establishment of transient glucose spikes

We first determined the window of time for a heightened level of blood glucose (**Figure 2A**). We varied the concentration of *D*-glucose that we injected from 0.25 g/kg to 2.0 g/kg. In all cases, injection of *D*-glucose results in a significant, transient increase in blood glucose concentration that peaks at 30 minutes and then returns to normal level after 120 minutes (**Figure 2B**). Thus, the laser injuries were performed 30 minutes after glucose injection. Further, an initial injection of 1.5 g/kg to 2.0 g/kg *D*-glucose leads to blood glucose concentration that exceed 300 mg/dl, the range of pathological hyperglycemia for non-fasting mice (King, 2012). We note that mice injected with *L*-glucose or with saline showed no significant changes in blood glucose levels (**Figure 2B**).

**Figure 2.**
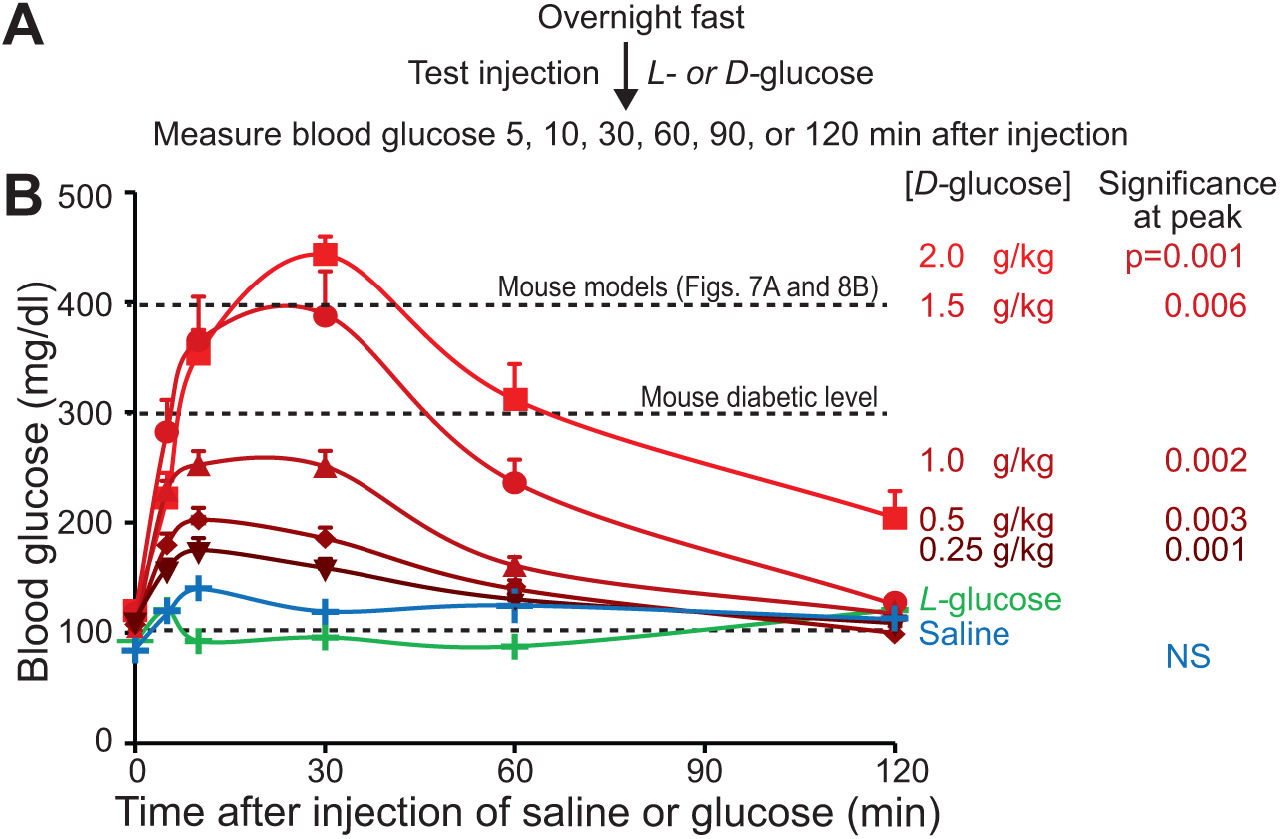
Blood glucose level measurements after injection of glucose into wild type mice. **(A)** Study design. **(B)** Data was taken after the injection of saline (control; 6 mice), *L*-glucose (inert control; 6 mice); or *D-*glucose (metabolic glucose) at concentrations of 0.25 g/kg (9 mice), 0.5 g/kg (7 mice), 1.0 g/kg (8 mice), 1.5 g/kg (5 mice), and 2.0 g/kg (9 mice); 50 mice total.

### Blinded study of vascular disruption from glucose spikes

Chronic hyperglycemia is associated with multiple morbidities that have been linked to vascular pathology. These include oxidative stress, inflammation, increased platelet reactivity, and endothelial dysfunction, any of which may result in a prothrombotic phenotype (Schneider, 2005). This raises the question of whether a high blood glucose level by itself is sufficient to induce susceptibility to thrombus formation in response to vascular tissue disruption. We addressed this question by examining the effects of targeted laser-induced disruption in normal, healthy mice subjected to experimental, transient hyperglycemia. In a fully blinded study, we used paired serial i.p. injections with an initial baseline dose of inert *L*-glucose, which cannot be metabolized and serves as osmotic control. This was followed by a second injection of either *L*-glucose or *D*-glucose; the nature of the second injection remained unknown until after data analysis of fluorescence changes was completed (**Figure 3A**).

**Figure 3.**
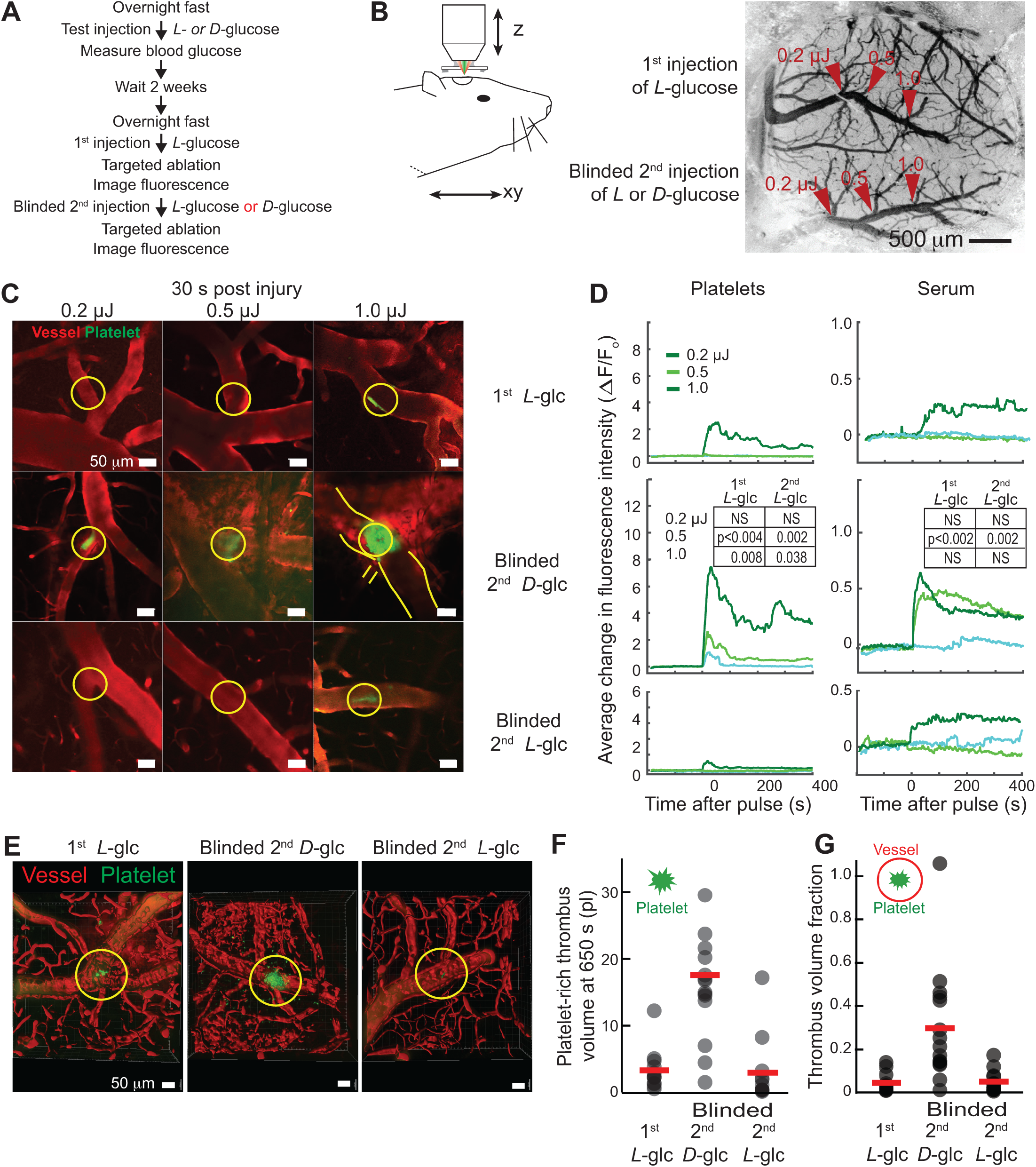
Double blinded study shows that transient hyperglycemia induced by injection of *D*-glucose but not *L*-glucose increases susceptibility to laser-induced vascular disruption. **(A)** Study design. **(B)** Schematic of set-up and image of surface vasculature of mouse cortex; arrow indicates branch of superior cerebral vein targeted with 0.2 µJ, 0.5 µJ, and 1.0 µJ laser energy, following treatment with 1^st^ injection of 2^nd^ injection of blinded *L-*glucose (upper vein) or *D-*glucose (lower vein). **(C)** Representative images of platelet (green) and serum (red) fluorescence in cortical veins targeted by 0.2 µJ, 0.5 µJ, and 1.0 µJ laser pulses, 30 s post laser damage, in wild type mice treated with a 1^st^ injection of *L-glucose* and a 2^nd^, blinded injection of *L-* or *D-*glucose. Site of focal damage is marked with yellow circle, along with the outline of the vein in selected images. **(D)** Effects of varying laser energy on the time course of the aggregation of platelets and blood extravasation induced by vascular disruption. Each curve is generated from the average change in vein fluorescence relative to baseline values. Each animal received a 1^st^ injection of *L-*glucose and a second injection of either *L-*glucose or *D-*glucose; 6 mice each for 12 mice total. A set of three veins per animal were irradiated 0.2 µJ, 0.5 µJ, and 1.0 µJ after the 1^st^ *L-*glucose injection and a second set was irradiated after the 2^nd^, blinded, injection. Data was smoothed with 2.4 s median filter. The insets give the Wilcoxon statistics for 1^st^ *L*-glucose and the blinded control 2^nd^ *L*-glucose relative to the experimental condition of the 2^nd^ *D*-glucose. **(E)** Three dimensional reconstructions of thrombi (green) within vasculature (red), collected 600 s post laser-induced vascular disruption. **(F)** Thrombi volumes at 600 s post laser-induced vascular disruption with 1.0 µJ irradiation. The red bar is the mean (3.3 ± 0.9 pl for 1st *L*-glc; 18 ± 3 pl for 2nd *D*-glc; 3.0 ± 1.4 pl for 2nd *L*-glc). For first injection with *L*-glucose the value does not differ from the baseline value of **Figure 1G** (p = 0.84). For second injection with *D*-glucose the value significantly differs from the baseline value (p < 7×10^−4^). For second injection with *L*-glucose the value does not differ from the baseline value (p = 0.75). **(G)** Thrombus volume fractions for 1.0 µJ irradiation; reanalysis of the data in panel F. The red bar is the mean (0.04 ± 0.01 for 1st *L*-glc; 0.30 ± 0.07 for 2nd *D*-glc; 0.05 ± 0.02 for 2nd *L*-glc). For the first injection with *L*-glucose the value does not differ from the baseline value of **Figure 1H** (p = 0.30). For a second injection with *D*-glucose the value significantly differs from the baseline value (p < 0.015). For a second injection with *L*-glucose the value does not differ from the baseline value (p = 0.45).

For each injection of *L-* or *D*-glucose, three pial venules were targeted for laser-induced disruption at energies of 0.2 µJ, 0.5 µJ, and 1.0 µJ each (**Figure 3B**). We used normalized changes in fluorescence to quantify the dynamics of platelet aggregation and serum extravasation. Platelet aggregation and serum extravasation were significantly greater after injections of *D*-glucose as compared to tissue damage observed after injection of inert *L*-glucose (**Figure 3C,D**). Volumetric reconstruction of the thrombi and the associated thrombus volume fractions also showed a greater value when formed in conjunction with *D*-glucose injections (**Figure 3E-G**). In the *D*-glucose injected mice the average thrombus volume fraction was 0.30 ± 0.07, as compared with a volume fraction of 0.04 ± 0.01 for the first *L*-glucose injection or a volume fraction of 0.05 ± 0.01 for the second *L*-glucose injection; the difference between *D*- and *L*-glucose is significant (p < 0.003 for 1st and p < 0.004 for 2nd *L*-glucose; **Figure 3G**). These data indicate that a transient elevation in blood glucose, by itself, is sufficient to render healthy mice significantly more susceptible to laser induced platelet aggregation and serum extravasation.

### Impact of sub-pathological as well as pathological blood glucose levels on susceptibility to disruption

To establish the susceptibility of vessels to laser-induced disruption, our double blind experiments (**Figure 3**) made use of a spike in blood glucose that attained levels normally seen in diabetic mouse models (**Figure 2**). We now address how blood glucose spikes of various magnitudes affect the cerebrovascular thrombo-inflammatory response. Healthy mice were subjected to serial i.p. injections with an initial baseline saline, followed by various concentrations of *D*-glucose that ranged from 0.25 g/kg, leading to a peak level of 170 mg/dl blood glucose (**Figure 2**) to 2.0 g/kg, leading to a peak level of 440 mg/dl blood glucose (**Figure 2**) (**Figure 4A**). Second injections of *L*-glucose and saline control served as controls (**Figure 4A**). For each pair of injections of saline*-* and *D*-glucose, three pial venules were targeted for laser disruption with 1.0 µJ laser pulses(**Figure 4B**). Platelet aggregation and serum extravasation were significantly greater than baseline after injections of *D*-glucose at all concentrations greater than 0.5 g/kg (**Figure 4C,D**); only injection of *D*-glucose at a concentration of 0.25 g/kg did not lead to significant, excessive damage (**Figure 4C,D**).

**Figure 4.**
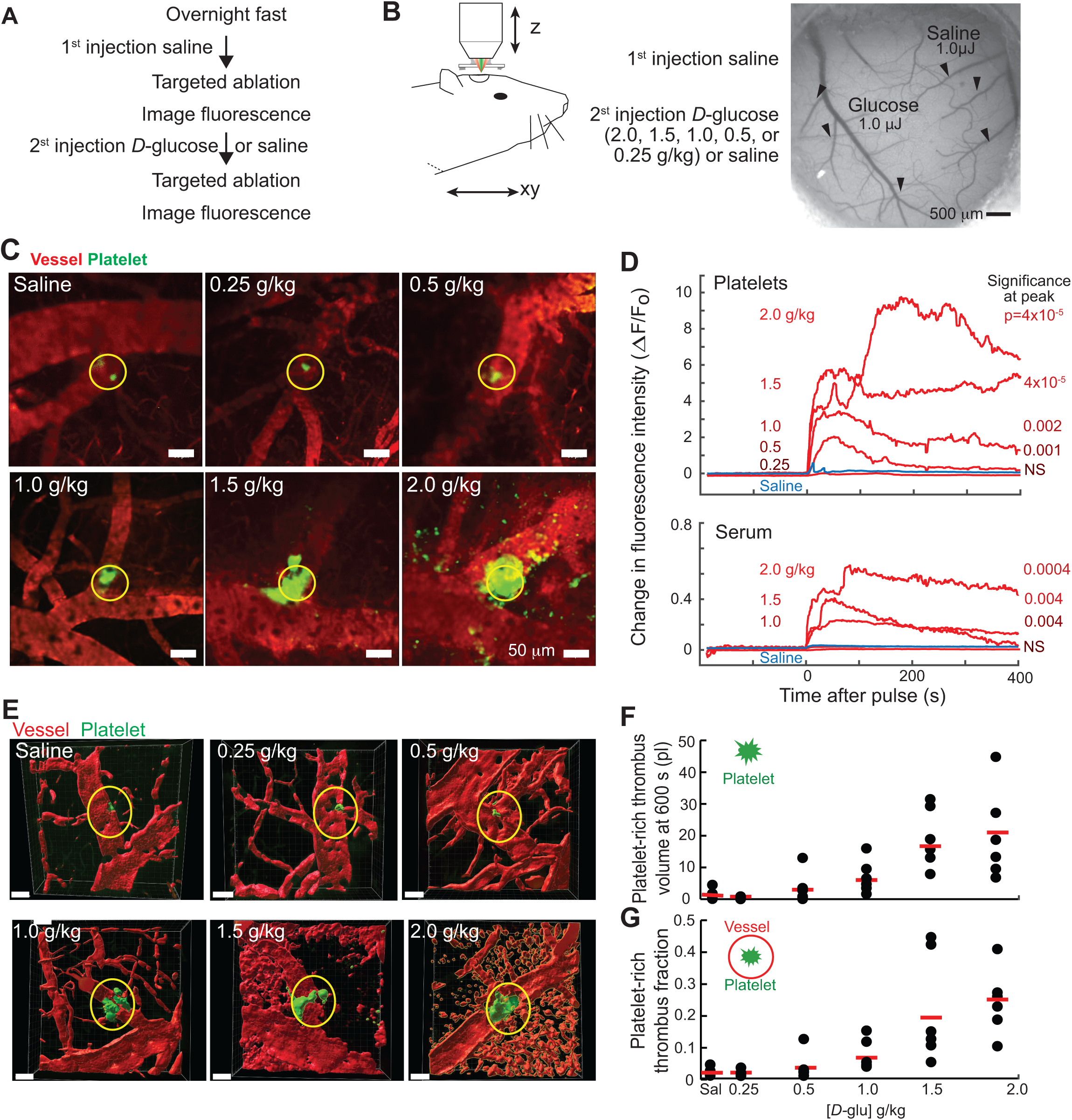
Normal, sub pathological and pathological blood glucose level effect on cerebrovascular susceptibility. **(A)** Study design. **(B)** Schematic of set-up and image of surface vasculature of mouse cortex; arrow indicates branch of superior cerebral vein targeted with 1.0 µJ laser energy, following treatment with 1^st^ injection of saline and *2*^nd^ injection of *D-*glucose, at 0.25, 0.5.1.0,1.5 or 2.0 g/kg, or a saline control. **(C)** Representative images of platelet (green) and serum (red) fluorescence in cortical veins targeted 1.0 µJ laser pulses, 30 s post laser-induced vascular disruption, in WT mice treated with a 1^st^ injection of saline and a 2^nd^ injection of *D-*glucose in concentrations of 0.25, 0.5.1.0,1.5 and 2.0 g/kg. Site of focal damage is marked with yellow circle, along with the outline of the vein in selected images. **(D)** Effects of 1.0 µJ laser energy on the time course of aggregation of platelets and blood extravasation. Each curve is generated from the average change in vein fluorescence relative to baseline values. Each animal received a 1^st^ injection of saline and a second injection of *D-*glucose. A set of three or four veins per animal for were irradiated 1at 1.0 µJ after the 1^st^ saline injection and after a second injection of 0.25, 0.5, 1.0, 1.5, or 2.0 g/kg D-glucose; 3 mice each for 15 mice total. As a control, a set of three veins per animal for were irradiated at 1.0 µJ after both a 1^st^ saline injection and a 2^nd^ saline; 12 mice total. The grand total is 27 mice. Data was smoothed with 2.4 s median filter. **(E)** Representative three dimensional reconstructions of thrombi (green) within vasculature (red), collected 600 s post laser-induced vascular disruption. **(F)** Thrombus volumes at 600 s post laser-induced vascular disruption. The red bar is the mean; only 7 of the 12 saline controls were used. For the injection with saline the value does not differ from the baseline value of **Figure 1G.** **(G)** Thrombus volume fractions; reanalysis of the data in panel G. The red bar is the mean.

Volumetric reconstruction of the platelet-rich thrombi and the associated thrombus volume fractions showed a greater value when formed in conjunction with *D*-glucose injections (**Figure 4E-F**). In the *D*-glucose injected mice the average thrombus volume fraction was significantly different compared to saline control for injections of 1.0 g/kg, (thrombus fraction 0.06 ± 0.02, p < 0.03), 1.5 g/kg (thrombus fraction 00.18 ± 0.06, p < 0.02), and 2.0 g/kg (thrombus fraction 0.24 ± 0.03, p < 3×10^−5^) (**Figure 4G**).

All told, these data (**Figures 2** and **4D,F,G**) indicate that a transient elevation in blood glucose that attains sub-pathological concentrations is sufficient to render healthy mice significantly more susceptible to laser induced platelet aggregation and serum extravasation.

### Diminution of damage by suppression of platelet activation

Thrombo-inflammation, as the immediate response to vascular tissue disruption, also includes neutrophil activation and recruitment (De Meyer et al., 2016). We therefore extended our analyses of the effects of transient hyperglycemia on vascular tissue disruption to specifically monitor the neutrophil responses in LysM-eGFP mice, in which blood myelomonocytic cells and in particular neutrophils are endogenously fluorescently labeled (Faust et al., 2000). Mice were successively split into four cohorts as a means to test the effect of blocking platelet mediated neutrophil activation (**Figure 5A**). In the first split, mice were i.p. injected with clopidogrel, an adenosine diphosphate receptor P2Y12 blocker that acts to suppress platelet hyperactivation, or they were injected with saline as a control. The injections were made the night before and the day of analysis. The second split was by an injection of either *L-* or *D*-glucose. We targeted a single pial venule per mouse with amplified laser pulses of 1.0 µJ (**Figure 5B**).

**Figure 5.**
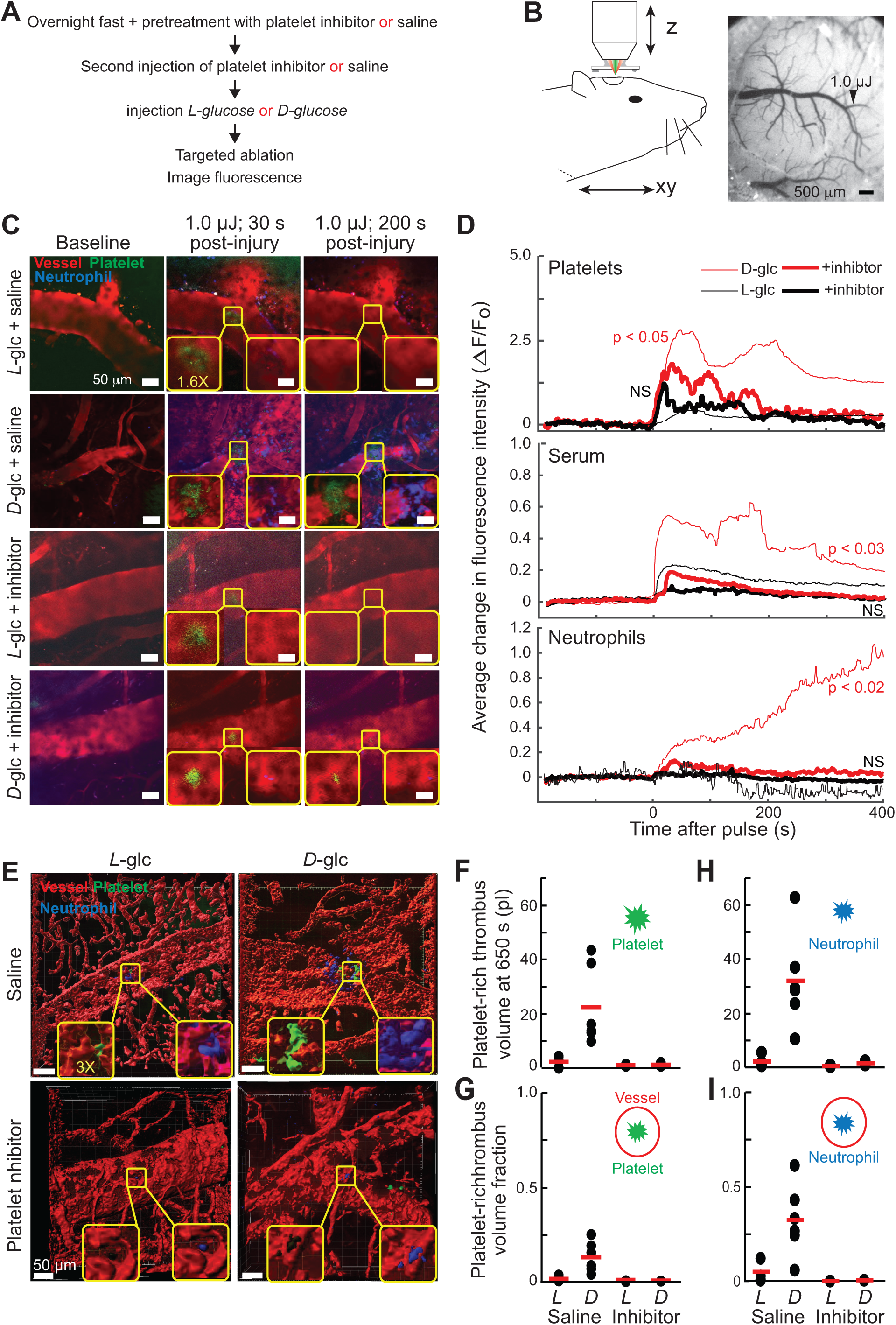
Transient hyperglycemia induced by *D*-glucose but not by *L*-glucose injection increases neutrophils recruitments following laser-induced vascular disruption. **(A)** Study design using lysM-eGFP mice with genetically labeled neutrophils. **(B)** Schematic of set-up and image of surface vasculature of mouse cortex; arrow indicates branch of superior cerebral vein targeted with 1.0 µJ laser energy, following pretreatment with platelet inhibitor or saline followed by injection of *L*-glucose or *D*-glucose. **(C)** Representative images of platelet (green), neutrophil (blue) and serum (red) fluorescence in cortical veins targeted with 1.0 µJ laser pulses, 0, 30 and 200 s post damage, in lysM-eGFP mice pretreated with saline, followed with injection *L*-glucose or *D*-glucose, or pretreated with platelet inhibitors (clopdogrel) followed by *L*- or *D*-glucose. Site of focal damage is marked with yellow circle, along with vein outline in some images. **(D)** Effects of 1.0 µJ laser energy on the time course of laser-induced thrombi, neutrophil recruitment and blood extravasation. Each curve is generated from the average change in vein fluorescence relative to baseline values. We use 6 mice and one vein per mouse in the saline pretreatment group, followed with injection *L*-glucose or *D*-glucose (6 mice per group or 12 total), and 4 mice and one vein per mouse in the platelet inhibitor group, also followed with injection *L*-glucose or *D*-glucose (4 mice per group or 8 total); grand total of 20 mice. Data was smoothed with 2.4 s median filter. P-values refer to Wilcoxon statistics for 1^st^ *L*-glucose and the blinded control 2^nd^ *L*-glucose relative to the experimental condition 2^nd^ *L*-glucose are computed for the initial platelet change and the steady-state serum extravasation. **(E)** Representative three dimensional reconstructions of thrombi (green) and neutrophil (blue) within vasculature (red), collected 600 s post laser-induced vascular disruption. **(F)** Thrombi volumes at 600 s post laser-induced vascular disruption. Same veins as in panel D (upper panel). The red bar is the mean (1.8 ± 0.7 pl for *L*-glc+saline; 22 ± 6 pl for *D*-glc+saline; 0.4 ± 0.1 pl for *L*-glc+inhibitor; 0.7 ± 0.2 pl for *D*-glc+inhibitor). For injection with *L*-glucose plus saline the values do not differ from the baseline values of **Figure 1G** (p = 0.47). For injection with *D*-glucose plus saline the thrombus volumes differ from the baseline value (p < 0.0008). For injection with *L*- or *D*-glucose plus inhibitor the values do not differ from baseline; p = 0.09 and p = 0.13, respectively. **(G)** Thrombus volume fractions; reanalysis of the data in panel F. The red bar is the mean (0.009 ± 0.003 for *L*-glc+saline; 0.13 ± 0.03 for *D*-glc+saline; 0.001 ± 0.001 for *L*-glc+inhibitor; 0.003 ± 0.001 for *D*-glc+inhibitor). For injection with either *L- or D-*glucose plus saline the value does not differ from the baseline values of **Figure 1H**; p = 0.70 and p = 0.15, respectively. For injection with *L*- or *D*-glucose plus inhibitor the values also do not differ from baseline; p = 0.15 and p = 0.16, respectively. The volume for of the platelet aggregates after injection with *D*-glucose is significantly greater than that with *D*-glucose + inhibitor (p < 0.031). **(H)** Neutrophil aggregate volumes at 600 s post laser-induced vascular disruption. Same veins as in panel D. The red bar is the mean (2.0 ± 0.7 pl for *L*-glc+saline; 32 ± 7 pl for *D*-glc+saline; ± 0.2 pl for *L*-glc+inhibitor; 1.5 ± 0.5 pl for *D*-glc+inhibitor). For injection with *D*-glucose plus saline the value significantly differs from baseline (p < 0.00020). For injection with *L*- or *D*-glucose plus inhibitor the values do not differ from baseline; p = 0.19 and p = 0.36, respectively. (**I**) Neutrophil aggregate volume fraction; reanalysis of the data in panel H. The red bar is the mean (0.05 ± 0.02 for *L*-glc+saline; 0.32 ± 0.08 for *D*-glc+saline; 0.001 ± 0.001 for *L*-glc+inhibitor; 0.005 ± 0.002 for *D*-glc+inhibitor). For injection with *D*-glucose plus saline the value significantly differs from baseline (p < 0.0018). For injection with *L*- or *D*-glucose plus inhibitor the values do not differ from baseline; p = 0.15 and p = 0.17, respectively.

In the absence of inhibitor, we observed that the *D*-glucose injected mice developed platelet aggregates that were significantly larger than those in *L*-glucose injected mice (**Figure 5C,D**). This is consistent with the previous, fully-blinded experimental result (**Figure 3D**). Neutrophil recruitment was also significantly greater in the *D*-glucose treated mice, with a linear growth of neutrophil binding (slope = 0.0023 ± 0.0001 s^-1^; **Figure 5D**). Critically, pretreatment with the platelet inhibitor clopidogrel significantly reduced platelet aggregation and neutrophil recruitment in both *L*- and *D*-glucose injected mice (**Figure 5C,D**). Pretreatment with clopidogrel also inhibited serum extravasation (**Figure 5C,D**). Volumetric reconstruction of platelet and neutrophil aggregation indicated significantly greater volumes in the *D*-glucose injected mice as compared to *L*-glucose injected mice and as compared to *D*-glucose injected mice with the platelet inhibitor (clopidogrel) (**Figure 5E,F**). In the *D*-glucose injected mice the platelet aggregates led to a thrombus volume fraction of 0.13 ± 0.03, significantly higher than in the *L*-glucose injected mice, albeit not higher than baseline control (**Figure 1H**). Similarly, the neutrophil aggregates occupied a thrombus volume fraction of 0.32 ± 0.08, which is significantly higher than in the *L*-glucose injected mice (p < 0.006); the relatively large volume results from the continued recruitment of neutrophils over the course of the measurements (Figure **5D**) and the large size of neutrophils relative to that of platelets. Platelet inhibitor treatment greatly diminished the aggregation of platelets for the *D*-glucose injected mice (**Figure 5F,G**) as well as aggregation of neutrophils (**Figure 5H,I)**, with the thrombus volume fraction reduced to 0.005 ± 0.002 (**Figure 5I**); both changes are significant (p < 0.03 and p < 0.02).

In total, we found that transient hyperglycemia was sufficient to render the cerebrovasculature more susceptible to vascular disruption, which is reflected not only in platelet aggregation and serum extravasation but also in neutrophils accumulating on top of platelet-rich thrombi. Under hyperglycemic conditions, hyperactivated platelets are not only an obstacle to homeostatic blood flow but also provide a nucleation point for neutrophils to ensue cerebrovascular inflammation.

### Diminution of damage by scavenging of reactive oxygen

A known complication of diabetes is the elevated production of reactive oxygen species (ROS) (Volpe et al., 2018). Excessive concentrations of ROS can hyperactivate platelets by depleting homeostatic vascular NO to form peroxynitrite(Forstermann et al., 2017). We asked if transient hyperglycemia, by itself, is a sufficient trigger for elevated production of ROS. To answer this, we made use of the mitochondria-targeted antioxidant Mitotempo, which is a ROS scavenger, and S-nitroso-N-acetyl-*DL*-penicillamine (SNAP), which is a nitric oxide (NO) donor. Mice were successively divided into six cohorts as a means to test the effect of scavengers (**Figure 6A**). In the first split, wild type mice were i.p. injected with Mitotempo, with SNAP, or with saline as a control. The second split was by an injection of either *L-* or *D*-glucose. We targeted a single pial vessel of each mouse with amplified laser pulses of 1.0 µJ (**Figure 6B**).

**Figure 6.**
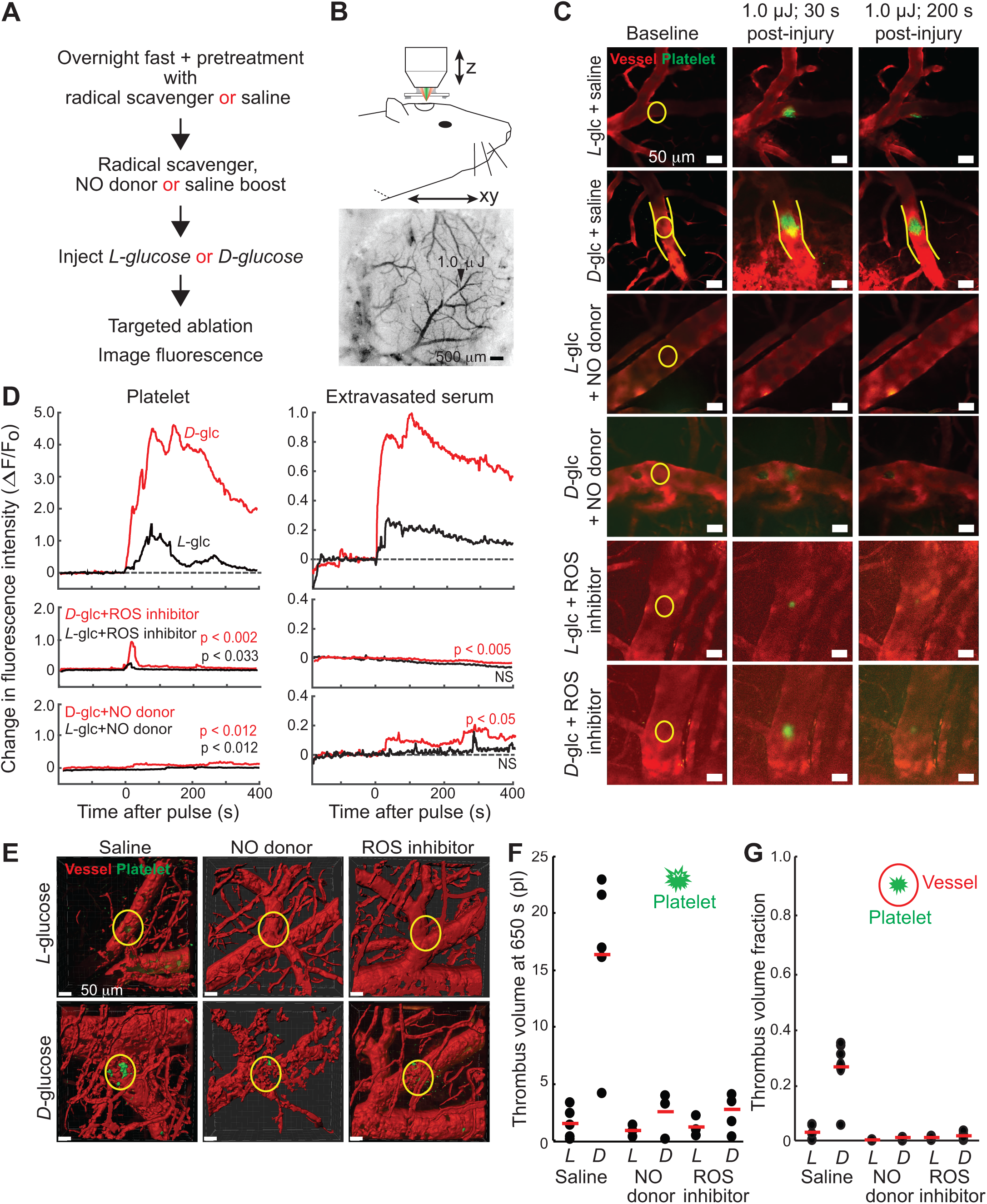
Involvement of hyperglycemia induced ROS and NO inhibition in susceptibility to laser-induced vascular disruption. **(A)** Study design. **(B)** Schematic of set-up and image of surface vasculature of mouse cortex; arrow indicates branch of superior cerebral vein targeted with 1.0 µJ laser energy, following pretreatment with NO donor, ROS scavenger or saline followed by injection of *L*-glucose or *D*-glucose. **(C)** Example images of platelet (green), and serum (red) fluorescence in cortical veins targeted with 1 µJ laser pulses, 0, 30 and 200 s post laser-induced vascular disruption in wild type mice pretreated and treated with saline, followed with injection *L*- or *D*-glucose, or pretreated and treated with ROS scavenger (MitoTempo) followed by *L*- or *D*-glucose, or treated with NO donor (SNAP) followed by *L*- or *D*-glucose. Site of focal damage is marked with yellow circle, along with vein outline in some images. **(D)** Effects of 1.0 µJ laser pulses on the time course of laser-induced thrombi, neutrophil recruitment and blood extravasation. Each curve is generated from the average change in vein fluorescence relative to baseline values. We use 12 mice and one vein per mouse irradiated at 1.0 µJ in the saline group (+ *L*- or *D*-glucose at 6 mice each). We use 8 mice and one vein per mouse irradiated at 1.0 µJ in the ROS scavenger group (+ *L*- or *D*-glucose at 4 mice each). We use 8 mice and one vein per mouse irradiated at 1.0 µJ in the NO donor group (+ *L*- or *D*-glucose at 4 mice each). The grand total is 28 mice. Data was smoothed with 2.4 s median filter. P-values refer to Wilcoxon statistics for treatments with ROS-inhibitor and NO-donor for *L*-glucose and *D*-glucose injected mice relative to untreated. We computed statistics for the initial platelet change and the steady-state serum extravasation. **(E)** Example three dimensional reconstructions of thrombi (green) within vasculature (red), collected 600 s post laser-induced vascular disruption. **(F)** Thrombus volumes at 600 s post laser-induced vascular disruption. The red bar is the mean (1.5 ± 0.6 pl for *L*-glc+saline; 16 ± 3 pl for *D*-glc+saline; 1.2 ± 0.5 pl for *L*-glc+NO donor; 2.8 ± 0.7 pl for *D*-glc+NO donor; 0.9 ± 0.3 pl for *L*-glc+ROS inhibitor; 2.6 ± 0.7 pl for *D*-glc+ROS inhibitor). For the injection of *L*-glucose the volume is consistent with baseline (**Figure 1G**; p = 0.25), for injection of D-glucose the volume differs from baseline (p < 0.0060). For injection of either *L*- or *D*-glucose + NO donor the volume is consistent with baseline (p = 0.11 and p = 0.62, respectively). For injection of either *L*- or *D*-glucose + ROS inhibitor the volume is consistent with baseline (p = 0.92 and p = 0.56, respectively). **(G)** Thrombus volume fractions; reanalysis of the data in panel F. The red bar is the mean (0.03 ± 0.01 for *L*-glc+saline; 0.26 ± 0.05 for *D*-glc+saline; 0.011 ± 0.004 for *L*-glc+NO donor; 0.016 ± 0.006 for *D*-glc+NO donor; 0.003 ± 0.001 for *L*-glc+ROS inhibitor; 0.010 ± 0.003 for *D*-glc+ROS inhibitor). For the injection of *L*-glucose the volume fraction is consistent with baseline (**Figure 1G**; p = 0.26), for injection of *D*-glucose the fraction differs from baseline (p < 0.0024). For injection of either *L*- or *D*-glucose + NO donor the fraction is consistent with baseline (p = 0.21 and p = 0.14, respectively). For injection of either *L*- or *D*-glucose + ROS inhibitor the volume is consistent with baseline (p = 0.16 and p = 0.10, respectively).

In the absence of antioxidants, we observed that the immediate acute response of *D*-glucose injected mice was an increase in platelet aggregation and serum extravasation after a laser pulse as compared to the effects observed in the *L*-glucose injected control mice (**Figure 6C,D**). Critically, pretreatment with Mitotempo or with SNAP reduced platelet aggregation and reduced serum extravasation in both *L-* and *D*-glucose injected mice (**Figure 6C,D**). Volumetric reconstruction of platelet aggregates in the *D*-glucose injected mice indicated that these volumes were significantly greater than platelet aggregate volumes in *L*-glucose injected control mice (**Figure 6E,F**). In the *D*-glucose injected mice the average platelet volume fraction after laser-induced disruption was 0.26 ± 0.05, significantly higher than for *L*-glucose injected control mice (p < 0.003) (**Figure 6G**). Injection of the ROS inhibitor Mitotempo dramatically and significantly reduced the thrombus volume fraction, to 0.010 ± 0.003, for the *D*-glucose injected mice (p < 0.002) (**Figure 6G**). These data are in agreement with past studies that connects ROS intermediates with cerebrovascular inflammation (Olmeza and Ozyurtbc, 2012). Further, injection of the NO donor SNAP significantly reduced the thrombus volume fraction to 0.016 ± 0.006 for the *D*-glucose injected mice (p < 0.002) (**Figure 6G**). In total, these data suggest a mechanism, in which excessive blood glucose fuels ROS production. This, in turns depletes, homeostatic NO and links hyperglycemia to cerebrovascular thrombus formation and inflammation.

### Models of chronic hyperglycemia

How does the venous response to laser-induced tissue disruption for transient hyperglycemic normal mice compare to that for established mouse disease models of diabetes? We investigated this for two mouse well known models. The first is a model of type 1 diabetes in which streptozotocin (STZ) injections have destroyed pancreatic β-cells (Wu and Huan, 2008) (**Figure 7**). In these mice with diabetes, fasting blood glucose levels are high, of order 400 mg/dl (**Figure 7A**). By comparison, in control, nondiabetic mice blood glucose levels are normal, of order 100 mg/dl (**Figure 7A**). The high blood glucose level in STZ mice could be transiently reduced with a single i.p. injection of insulin (1 U/kg equal to ∼ 120 mg/dl) (**Figure 7A**). To study pial venule disruption during hyperglycemia and insulin normalization of hyperglycemia, we compared responses to paired injections of saline-saline or saline-insulin, performed serially in individual mice (**Figure 7B**). We used changes in fluorescence to quantify the dynamics of platelets aggregation and serum extravasation in response to laser injuries at 0.2 µJ, 0.5 µJ, and 1.0 µJ (**Figure 7C**). Both platelet aggregation and serum extravasation were observable at the 1.0 µJ threshold level for disruption after the first saline injection and in cases with a second saline injection (**Figure 7D,E**). However, injection with insulin significantly reduced irradiation triggered platelet aggregation and abrogated serum extravasation (**Figure 7D,E**). Laser-induced tissue disruption produced platelet aggregates whose volume and volume fractions were significantly reduced, to near control levels, during insulin treatment (middle row, **Figure 7D,E**), when glucose levels are reduced to near normal levels (**Figure 7A)**. A similar reduction in thrombus volume during insulin treatment was observed (**Figure 7F,G**). This effect is substantial; the average thrombus volume fraction was over three-times as large in STZ mice (0.22 ± 0.04, mean ± SE; **Figure 7H**) as compared to normoglycemic mice (0.072 ± 0.024; **Figure 1H**) and insulin controls (0.035 ± 0.005; **Figure 7H**); the thrombus volume increase in STZ mice was statistically significant (t-test, p < 0.01).

**Figure 7.**
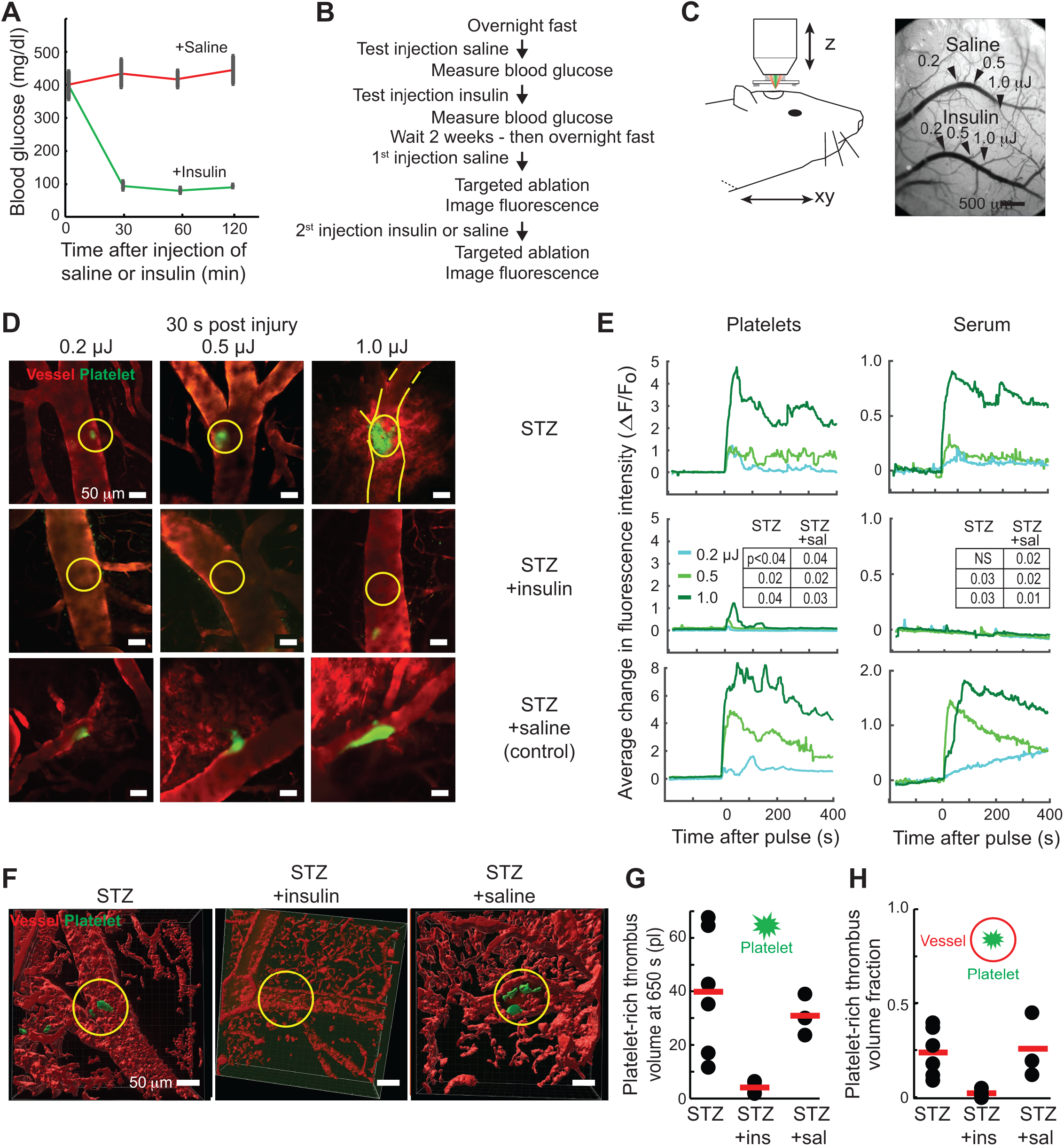
STZ induced hyperglycemia increases susceptibility to laser-induced vascular disruption. **(A)** Blood glucose level measurements (6 mice) over 120 minutes after injection of saline (control) in STZ mice followed by use of the same animals after injection of insulin. The bars are the SEM. **(B)** Study design. **(C)** Schematic of set-up and image of surface vasculature of mouse cortex; arrow indicates branch of superior cerebral vein targeted with 0.2 µJ, 0.5 µJ, and 1.0 µJ laser energy, following treatment with saline (upper vein) or insulin (lower vein). **(D)** Representative images of platelet (green) and serum (red) fluorescence in cortical veins targeted by 0.2 µJ, 0.5 µJ, and 1.0 µJ laser pulses, 30 s post laser-induced vascular disruption, in STZ mice treated with saline and STZ mice treated with insulin. Site of focal damage is marked with yellow circle, along with vein outline in some images. **(E)** Effects of varying laser energy on the time course of laser-induced thrombi and blood extravasation. Each curve is generated from the average change in vein fluorescence relative to baseline values. We used 9 mice total. All 9 mice received a 1^st^ injection of saline and each of 3 veins in each animal were irradiated at 0.2 µJ, 0.5 µJ, and 1.0 µJ. Then 6 of the 9 mice received a 2^nd^ injection of insulin (1 unit/kg) and each of 3 additional veins in each animal were irradiated at 0.2 µJ, 0.5 µJ, and 1.0 µJ. Further, 3 of the 9 mice received a 2^nd^ injection of saline and each of 3 additional veins in each animal were irradiated at 0.2 µJ, 0.5 µJ, and 1.0 µJ. Data was smoothed with 2.4 s median filter. We determined the effect of insulin suppression of susceptibility to laser-induced damage for the first 100 s after the pulse by calculating the rank order (one sided non-paired Wilcoxon) for STZ + insulin relative to STZ alone for each energy level as well as the ratio for STZ + insulin relative to STZ + saline for each energy level. NS - not significant or p > 0.05. **(F)** Representative three dimensional reconstructions of thrombi (green) within vasculature (red), collected 600 s post laser-induced vascular disruption. **(G)** Thrombi volumes at 600 s post laser-induced vascular disruption. Same veins as in panel D. The red bar is the mean (40 ± 9 pl for STZ; 3.6 ± 0.7 pl for STZ+insulin; 32 ± 4 pl for STZ+saline). For STZ only the value significantly differs from the baseline value of **Figure 1G** (p < 1.1×10^−4^). For STZ + insulin the value does not differ from the baseline value (p = 0.97). For STZ + saline the value significantly differs from the baseline value (p < 1.3×10^−5^) and is equivalent to the STZ only value (p = 0.55). **(H)** Thrombus volume fractions; reanalysis of the data in panel F. The red bar is the mean (0.22 ± 0.05 for STZ; 0.035 ± 0.006 for STZ+insulin; 0.24 ± 0.09 for STZ+saline). For STZ only the value significantly differs from the baseline value of **Figure 1H** (p < 0.01). For STZ + insulin the value does not differ from the baseline value (p = 0.27). For STZ + saline the value significantly differs from the baseline value (p < 0.02) and is equivalent to the STZ only value (p = 0.80).

To model type 2 diabetes, we made use of leptin-receptor knock-out *db*/*db* mice. These mice are significantly overweight (**Figure 8A**) and display chronic high blood glucose levels (∼ 320 mg/dl) as compared to control mice (∼ 110 mg/dl) (**Figure 8B**). This model of hyperglycemia cannot be reversed with insulin (Chen et al., 1996), which indicates insulin resistance. The effects of laser-induced disruption to venules, at pulse energies of 1.0 µJ, were compared between *db*/*db* hyperglycemic versus control mice with normal blood glucose (**Figure 8C,D**). The dynamic responses in these two groups demonstrated that there were significantly larger platelet aggregations and serum extravasation in the *db*/*db* mice, as compared to normoglycemic mice (**Figure 8E,F**). Volumetric reconstructions of platelet aggregates after laser-induced tissue disruption showed that obesity related hyperglycemia resulted in significantly greater aggregate volumes in *db/db* mice relative to control mice (**Figure 8G,H**). The thrombus volume fractions of platelet aggregates in *db/db* mice were 0.25 ± 0.08, which is significantly larger (p < 0.02) than the thrombus volume fractions for control normoglycemic mice (**Figures 1H** and **8I**).

**Figure 8.**
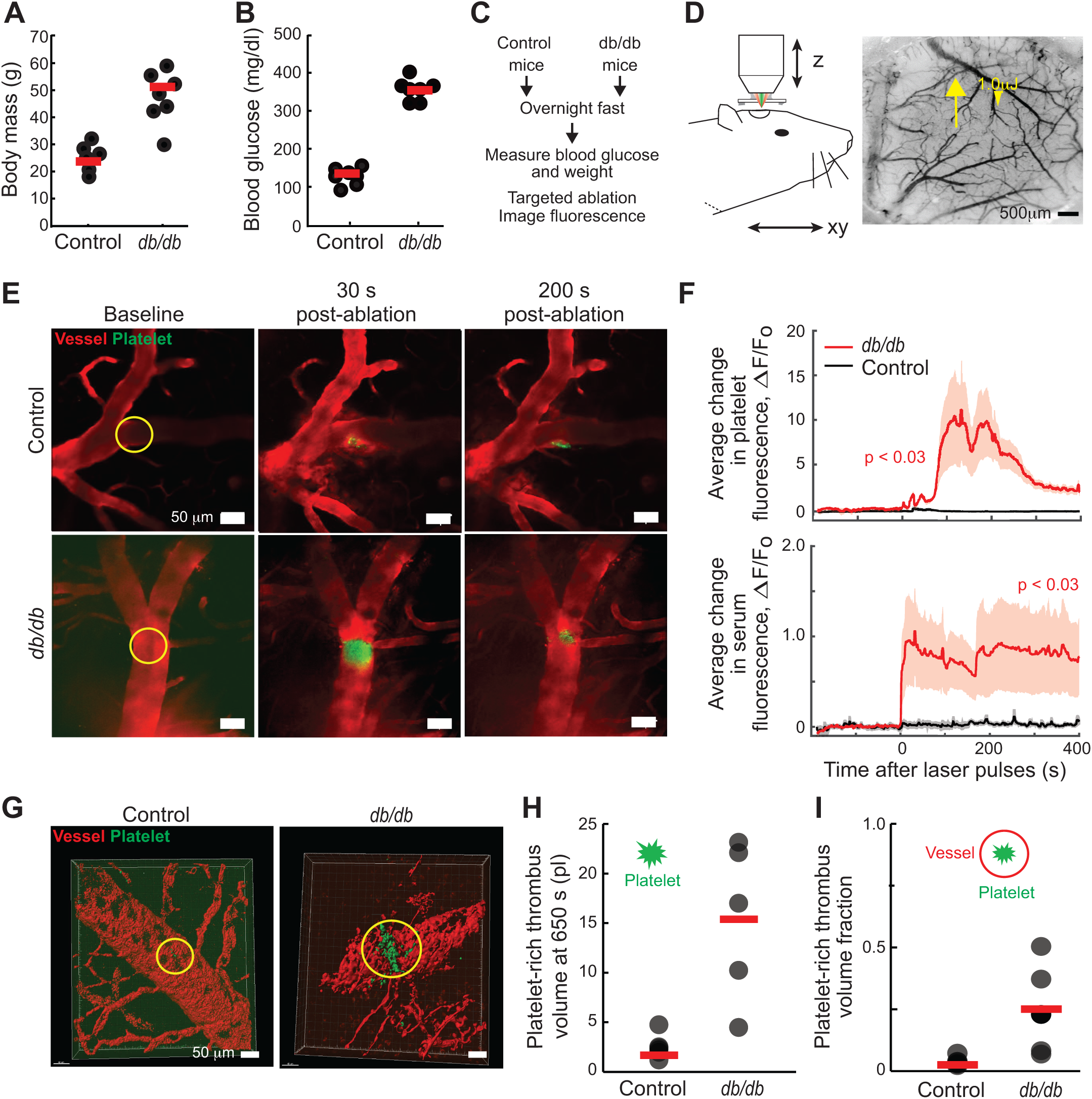
Obesity related type 2 hyperglycemia increases susceptibility to vascular disruption. **(A)** Body mass measurements in *db/db* in compare to wild type control mice (8 *db/db* and 8 WT mice); the differences are significant with p < 10^−12^. The red bar is the mean; 49.8 ± 1.0 g for *db/db* (mean ± SD) for 25.8 ± 0.5 g for WT. **(B)** Blood glucose level measurements in *db/db* in compare to wild type control mice; the differences are significant with p < 10^−7^. The red bar is the mean (137 ± 9 mg/dl for control; 351 ± 24 mg/dl for *db/db*). Same mice as in panel A. **(A)** Study design. **(D)** Schematic of set-up and image of surface vasculature of mouse cortex; arrow indicates branch of superior cerebral vein targeted with 1.0 µJ laser energy. **(E)** Representative fluorescent images of platelet (green) and serum (red) fluorescence in cortical surface veins, before and 30 s and 200 s post laser-induced vascular disruption using 1.0 µJ laser pulses, in wild type control mice in comparison to *db/db* mice. Focal region of the excitation spot is highlighted by the yellow circle. **(F)** Effects of 1.0 µJ laser pulses on the time course of laser-induced platelet-rich thrombi and serum extravasation. Each curve is generated from the average change in vein fluorescence relative to baseline values. We used 5 *db/db* mice with one vein targeted per mouse and 6 WT mice with one vein targeted per mouse, for a grand total of 11 mice. Data was smoothed with 2.4 s median filter; the light red bands are the 1 SD intervals. The changes for *db/db* mice are significant relative to control animals for both platelets (one-sided Wilcoxon for initial 100 s after pulse) and serum extravasation (near steady-state, i.e., 300 - 400 s after pulse). **(G)** Representative three dimensional reconstructions of thrombi (green) within vasculature (red), collected 600 s post laser-induced vascular disruption. **(H)** Thrombi volumes at 600 s post laser-induced vascular disruption. Same veins as in panel D. The red bar is the mean (1.6 ± 0.6 pl for control; 15 ± 4 pl for *db/db*). For control mice the value does not differ from the baseline value of **Figure 1G** (p = 0.17). For *db/db* mice the value is significantly different from the baseline value (p < 0.0016). **(I)** Thrombus volume fractions; reanalysis of the data in panel F. The red bar is the mean (0.030 ± 0.008 for control; 0.25 ± 0.08 for *db/db*). For control mice the value does not differ from the baseline value of **Figure 1H** (p = 0.16). For *db/db* mice the value is significantly different from the baseline value (p < 0.019).

We conclude that chronic hyperglycemia triggered by insulin deficiency (STZ type 1 diabetic model; **Figure 7**) or by insulin resistance (obese, *db*/*db* mice; **Figure 8**) rendered murine vessels more susceptible to enhanced thrombus formation in response to laser-induced cerebrovascular tissue disruption. Moreover, in both chronic hyperglycemia mouse models, a thrombus volume fraction of 0.2 or more is occupied by an aggregate of platelets; this presents obvious threats to normal blood flow.

### Meta-analysis of normoglycemia and transient hyperglycemia results

We purposely conducted separate controls for each set of experimental conditions as a means to minimize potential systematic variability between experiments, such as may occur with different litters of mice despite matching for age and sex. This yielded multiple sets of control experiments that involved the same or nearly the same conditions. We now considered a meta-analysis across experimental groups based on a one-way ANOVA. For the control condition of solely wild type animals (**Figure 1**) or wild type plus saline injection (**Figures 4 and 8**), we could not reject the null hypothesis, i.e., their mean values were not significantly different, yielding a thrombus volume fraction of 0.041 ± 0.011 (25 mice). For the control condition of wild type animals injected with *L*-glucose (**Figures 3, 5, and 6**), we found that the combined results did not differ significantly from each other, yielding a thrombus volume fraction of 0.035 ± 0.009 (23 mice). In fact, the data across all control groups do not differ significantly from each other and yield a grand thrombus volume fraction of 0.038 ± 0.007 (48 mice).

A similar meta-analysis was performed for all wild type animals injected with 2.0 g/kg of *D*-glucose (**Figures 3-6**), but no subsequent pharmacological agents. We found a thrombus volume fraction of 0.24 ± 0.04 (29 mice), i.e., six-times the size of the thrombus under control conditions and significantly different (p < 3 × 10^−9^).

## DISCUSSION

Our experimental approach allowed us to test the effect of a lone, transient spike in blood glucose independently from diabetes comorbidities. The essential data is based on a fully-blinded experimental design and reveals that a pulse of high blood glucose (**Figure 2**) administered to normal healthy animals, but not a pulse of an administered inert analogue, is sufficient to dramatically increase the platelet-rich thrombotic response to laser-induced vascular tissue disruption (**Figure 3**) This suggests that the susceptibility to vascular thrombo-inflammation is heightened by transient hyperglycemia, even at prediabetic levels of blood glucose (**Figure 4**). The extent of thrombi formation rivels that for the diabetes models (**Figures 7** and **8**), while the extent of extravasated serum is well above control levels but less than that for the type 1 and type 2 diabetes models (**Figure 9**). Lastly, a transient spike in blood glucose not only aggravates laser-induced platelet aggregation, but also intensifies neutrophil accumulation at the lesion site (**Figure 5**).

**Figure 9.**
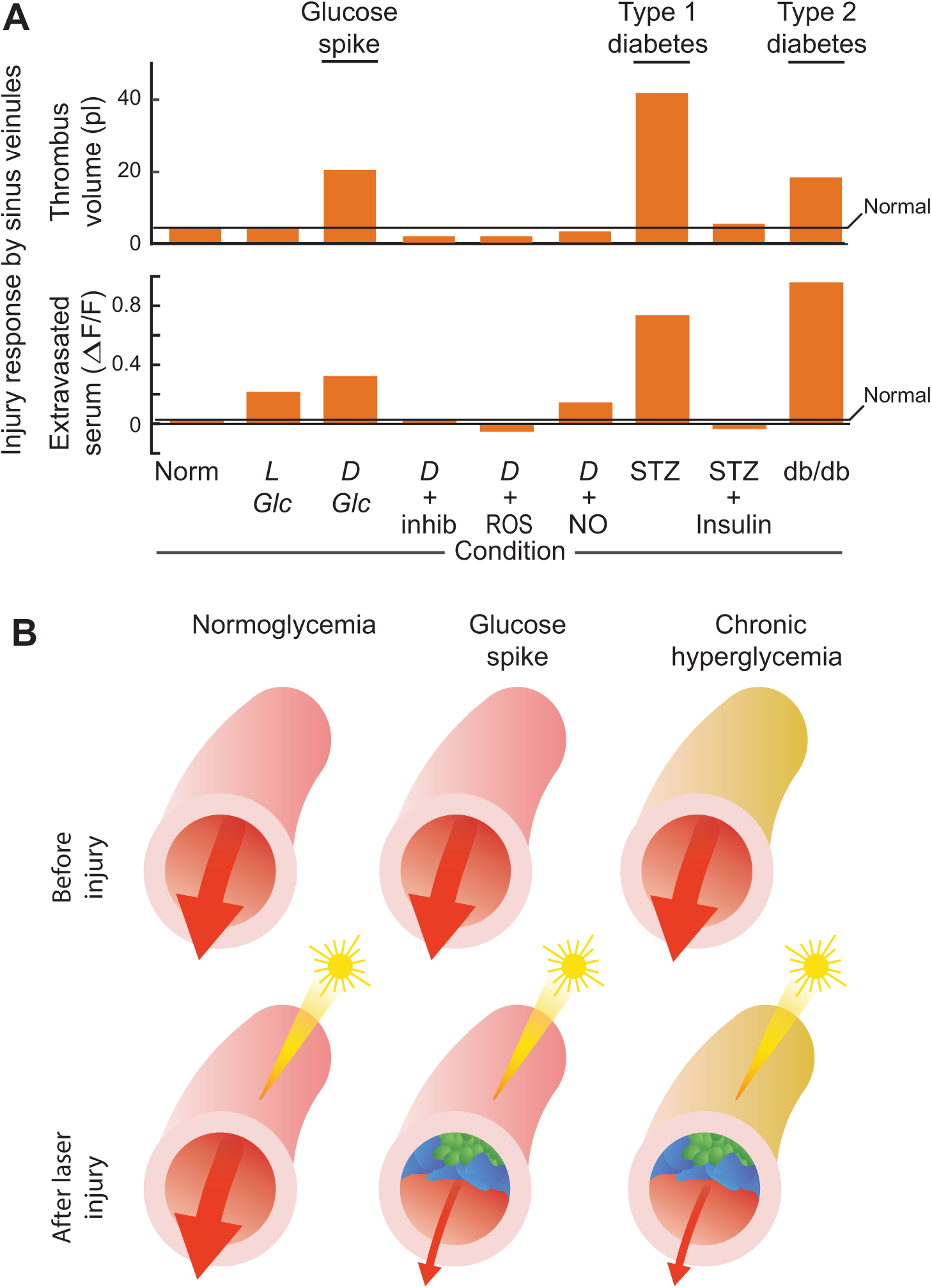
Transient vs. chronic hyperglycemia. **(A)** Combined effects of 1.0 µJ ultra-short pulsed laser energy on the peak of laser-induced of platelet fluorescence and the peak of laser-induced of serum extravasation fluorescence. **(B)** Graphical summary.

Our control studies tested the consequence of insulin deficiency, i.e., type 1 diabetes (STZ) (**Figure 6**). Delivery of minimally energetic laser pulses induces significant cerebrovascular blood thrombi in mice with diabetes, while having only marginal effect in control mice (**Figures 1 and 6**). The specificity of hyperglycemia is supported by the blunting of the effects of laser-induced disruption on platelet hyper-responsiveness when normal blood glucose levels are transiently restored by the injection of insulin (**Figure 6**). Note that insulin may also act as an antithrombic and anti-inflammatory agent independently of regulation of blood glucose (Patti et al., 2019). Diabetes from insulin resistance, i.e., in obesity-related type 2 diabetes (db/db) (**Figure 7**) also resulted in increased susceptibility to laser-targeted vessel disruption. The use of db/db mice represents a challenge, as they are hyper-insulinemic and cannot be treated with supplementary insulin (Genuth et al., 1971). Nonetheless, our findings suggest that conditions in which hyperglycemia is accompanied by hyperinsulinemia similarly render mice more susceptible to thrombo-inflammation (**Figure 8**).

A body of work on the interactions between the chronic hyperglycemic state and cerebro-vascular inflammation have used classic stroke models (Chen et al., 2019). Cerebrovascular stroke as modeled by transient occlusion of the middle cerebral artery occlusion in rodents (Zea Longa et al., 1989) results in larger infarcts at subacute time points, i.e., less than 24 hours, under hyperglycemic as compared to normoglycemic conditions (Denorme et al., 2021; MacDougall and Muir, 2011). Photothrombotic disruption of the vascular lumen (Schaffer et al., 2006), which models microvascular occlusions, results in worse outcomes in STZ diabetic mice (Watson et al., 1985), as compared to normoglycemic mice (Tennant and Brown, 2013). However, interpretation of hyperglycemic effects from photothrombotic vessel damage may be confounded by the production of injurious ROS that is stimulated by the photothrombotic agents (Dietrich et al., 1987).

What is the mechanism of heightened sensitivity to vascular damage in hyperglycemia? In vitro studies have been critical in suggesting that elevation in glucose levels promotes a variety of pro-coagulation events across vascular cell types. Blood vessel endothelial cells are stimulated by high glucose to increase their production of ROS (Nishikawa et al., 2000), which in turn decreases bioavailability of NO, a coagulation cascade inhibitor (Guzik et al., 2002). Elevated glucose activates constituent blood cell types including isolated platelets (Bridges et al., 1965), neutrophils and monocytes (van Oostrom et al., 2004) release of pro-coagulant intermediates (Kraakman et al., 2001) and/or up-regulation of cell surface molecules that promote cell adhesion, platelet-rich thrombi (Colwell and Nesto, 2003; Zhu et al., 2012), as well as neutrophil recruitment (Clark et al., 2007; Jenne et al., 2013). Consistent with the in vitro studies, we show that pretreatment of mice with MitoTempo, a potent scavenger of mitochondrial superoxide, as well as pretreatment with the NO donor SNAP, mitigate hyperglycemia-induced susceptibility to laser-induced thrombosis and serum extravasation (**Figures 6** and **9**).

The current study, combined with past *in vitro* studies (Nishikawa et al., 2000) supports a hypothesis that hyperglycemia, whether it is chronic or transient, can promote exacerbation of platelet thrombus formation in response vascular tissue disruption. Therapies to reduce ROS production and to increase NO levels may be sufficient to minimize thrombotic damage during transient hyperglycemia (**Figure 9**). A pragmatic means to minimize ROS vascular damage may involve dietary strategies that minimize fuel for mitochondrial ROS and maximize nutritional antioxidants (Mafra et al., 2021). This suggests that dietary lowering of blood glucose in the context of sickness behavior lethargy and food avidness can thus prove protective against thrombotic inflammation. These notions agree with the finding by Medzhitov and colleagues (Wang et al., 2016). that fasting that leads to hypoglycemia is protective from bacterial inflammation (Wang et al 2016). Lastly, studies support the clinical need to prevent glycemic variation, i.e., spikes in blood glucose levels in individuals with known risk factors for thrombosis (Ceriello et al., 2019). These risk factors include aging, heart failure, chemotherapy, and prolonged bed rest (Anderson and Spencer, 2003; Crane et al., 2013), as well as SARS-CoV-2 infection (Lim et al., 2021; Mazucanti and Egan, 2020), which increases susceptibility to vascular disease (Aid et al., 2020) and large-vessel stroke (Oxley et al., 2020).

## Acknowledgments

We thank Marta Koch, Dorian McGavern, Helena Shaked, Orian Shirihai and Bart Weijts for discussions, Celine Mateo for assistance with data analysis, Rodolfo Figueroa-Mercado for assistance with the blood glucose measurements, and Hannah Liechty for assistance with the fully blinded experimental design. This work was sponsored by National Institutes of Health grants R01 NS108472 and R35 NS097265. Image analysis was performed at the UCSD School of Medicine Microscopy Core, funded by National Institutes of Health grant P30 NS047101.

## Author Contributions

DK and IS conceived the study. AD and DK secured funding. YC and XJ analyzed pial vasculature. CF, DK and RL constructed the in vivo experimental set-up. IS performed the in vivo experiments with the assistance of TB and PM. CF performed all statistical tests. BF, DK, IS and PS discussed and organized the results. BF, DK and IS wrote the manuscript. DK attended to the myriad of university rules and forms that govern environmental health and safety, including the ethical use of animals as well as the use of chemicals, controlled substances, hazardous substances, and lasers.

## METHODS

### EXPERIMENTAL MODEL AND SUBJECT DETAILS

All animal procedures were approved by the Institutional Animal Care and Use Committee at the University of California, San Diego (UCSD) and were performed under the guidance of the UC San Diego Center for Animal Resources and Education. Our subjects are summarized in **Table 1**.

**Table 1.**

| Strain | Gender | Age (weeks) | Objective | Treatment | Mice / treatment | Veins / mouse | Total veins |
| --- | --- | --- | --- | --- | --- | --- | --- |
| B6 WT | male | 8-12 | Control | - | 40 | 4-7 | 220 |
| B6 WT (STZ) | male | 8-12 | T1DM | STZ ± I.P. insulin | 10 | 6 | 60 |
| B6 WT | male | 8-12 | Transient hyperglycemia | I.P. *L/D*-glc or saline | 50 (Fig. 2) | - | - |
|  |  |  |  |  | 12 (Fig. 3) | 3 | 36 |
|  |  |  |  |  | 27 (Fig. 4) | 3-4 | 95 |
| *db/db* | male | 8-12 | T2DM obesity | - | 9 (5 imaged) | 1 | 5 |
| B6 WT | male | 8-12 | Control to T2DM | - | 8 | 1 | 8 |
| LyZ2 | male | 8-12 | Imaging neutrophils | I.P. ± anticoagulants | 18 | 1 | 18 |
| B6 WT | male | 8-12 | ROS/NO | I.P. *L/D*-glc ± inhibitor/donor | 26 | 1 | 26 |

WT = wild type.
We supply data from 200 mice in total.

## METHOD DETAILS

### Chronic hyperglycemia

We used streptozotocin (STZ) induced type 1 diabetes mice. STZ induced mice display chronic hypoinsulinemia and hyperglycemia. This chronic hyperglycemia can transiently reverse by a single i.p. injection of insulin (1 unit/kg (TOCRIS-#3435). STZ treated mice were purchased from The Jackson Laboratory (**Table 1**). As a type 2 diabetes mellitus, we used leptin receptor deficient *db/db* mice The Jackson Laboratory (**Table 1**), which display obesity and insulin resistance evidenced by sever hyperglycemia.

### Transient hyperglycemia

To transiently modulate blood sugar, *L*-glucose or *D*-glucose (Sigma) solutions were i.p. injected at 2 gm/kg mouse body weight, 20 % (w/v) in saline, following 18 hours fast, 30 minutes prior to the laser induced vascular disruption. Blood sugar levels were monitored (3-5 µl samples) from the tail vein with a glucometer (Contour®).

### IP injections

Pharmacological modulators were administered by i.p. injection:

1. The ADP receptor P2Y12 blocker, Clopidogrel (1 mg/kg) (TOCRIS #2490) 1 day before imaging.
2. The NO donor S-nitroso-N-acetyl-DL-penicillamine (SNAP) (Sigma-Aldrich #N3398) (5mg/kg) was administrate 1 hour before imaging.
3. The mitochondrial oxygen free radical scavenger, MitoTempo (10 mg/kg) (Sigma SML 0737) was administrated 1 day and a second boost 1 hour before imaging.

### Surgery

Eight-week-old mice were anesthetized with isoflurane, 3 % (v/v) oxygen for induction and1.5 % – 2.5 % (v/v) for maintenance, from a precision vaporizer. Reflexes and breathing were visually monitored through the entire surgical procedure to ensure a deep plane of anesthesia. Body temperature was maintained at 37°C with a heating pad with feedback regulation (FHC, model 40-90-8D). The animal was then placed in a stereotaxic frame, and the periosteum on the parietal and occipital plates was exposed, only skull sutures were covered with low viscosity cyanoacrylate glue (Loctite, no. 4104) to reinforce stability between skull plates, and a 1.5 mm by 1.5 mm region of skull over somatosensory cortex was thinned with a 250 μm drill bur coupled to a low vibration drill (Osada, EXL-M40) to form a transcranial window (Drew et al., 2010; Shih et al., 2012). The thinned bone was dried and covered with cyanoacrylate glue (Loctite, no. 401) and a number 0 glass coverslip and a T-shaped metal implant was glued onto the skull for head-fixation. The remaining exposed bone and the implant were covered with cyanoacrylate glue and dental cement (Grip Cement, Denstply no. 675571) to increase stability. Buprenorphine hydrochloride (Buprenex, Reckitt Benckiser Pharmaceuticals) was provided subcutaneously for analgesia (7 μg) as the animal recovered from surgery. Mice were allowed to recover for 48-72 hours before experiments were performed.

### Retro-orbital injection

Labeling of serum was achieved with Texas red dextran (2 MDa) (Thermo-Fisher T6134) or Alexa Fluor 405 (AF405) dextran (2 MDa). Labelling of platelets was achieved using Biotin anti-mouse CD41 (integrin alpha 2b) antibody (BioLegend #133930) conjugated with Strepavidin Alexa Fluor™ 594 or Strepavidin 405 (ThermoFisher Scientific #S11227, #S32351).

### In vivo imaging and laser induction of intravascular blood platelet aggregates

Live images of mouse cerebral vasculature were obtained with a system of local design in which a two-photon laser scanning microscope incorporates a beam line from an amplified 100-femtosecond pulsed laser (Tsai and Kleinfeld, 2009) (**Figure 1C**). Incarnations of this versatile system has been used in our past work on precision tissue ablation for histology (Tsai et al., 2005), the induction of microstokes in vessels deep to the cortical surface (Nishimura et al., 2006), and for precision osteotomies. We extend it to produce micrometer-sized regions of damage to a vessel lumen.

#### Cojoining of beams

The beam line for imaging by two-photon microscopy utilizes low-energy, 100-fs pulses at a repetition rate of 76-MHz and a center wavelength of 840 nm. This beam is generated by a commercial fiber laser oscillator (Chameleon Discovery; Coherent Inc.). It is naturally polarized. The beam line for laser induction of an intravascular thrombus by targeted irradiation utilizes high-energy, 100-fs pulses at a repetition rate of 1-kHz and a center wavelength of 805 nm. This beam is generated a commercial combined seed laser and multi-pass amplifier (Astrella; Coherent Inc.). This beam too is naturally polarized. The imaging beam and the ablation beam were combined before the microscope objective with a polarizing beam splitter. A half-wave plate (λ/2) for each beam rotates the polarization of the amplified pulses to lie vertical and that of the laser oscillator to lie horizontal, so that both beams are perpendicular to each other yet lie of different polarization axes of the polarizing beam splitter (**Figure 1C**). These beams pass through the objective;. we used a 0.95-NA, 25X, water-dipping objective (Leica).

#### Cofocusing of beams

We centered the ablation beam within the area that is raster-scanned by the imaging beam so that damage occurred at the center of the imaging field. We further focused the two beams to the same focal plane by slight alteration of the vergence of the ablation beam (**Figure 1C**); e.g., convergence results in an increase in the depth of focus and vice versa.

#### Gating and energy control

The energy per pulse of the ablation beam was attenuated with successive sets of half-wave plates, polarizing beam splitters, and beam dumps. The number of pulses was controlled by a mechanical shutter (Uniblitz LS3Z2 shutter and VMM-D1 driver; Vincent).

#### Imaging

Individual frames were acquired at 2.13 Hz using standard galvanometric scanners (**Figure 1C**). We detected fluorescence with one of two schemes. For labeling of blood serum and platelets, we imaged Alexa Fluor™ 405 to monitor blood flow and serum extravasation (emission filtered at 457 ± 25 nm) and imaged Alexa Fluor™ 594 to monitor platelets (emission filtered at 615 ± 20 nm). For labeling of blood serum, platelets, and neutrophils, we imaged Alexa Fluor™ 405 to monitor platelets (emission filtered at 457 ± 25 nm), Texas Red to monitor blood flow and serum extravasation (emission filtered at 615 ± 20 nm), and EGFP to monitor neutrophils (emission filtered at 512 ± 12 nm).

We imaged single frames at the focal plane of the laser-induced damage for the first 412 s after laser-induced vascular disruption and then performed a volumetric reconstruction between 413 and 882 s after the laser-induced vascular disruption. The scan was arranged to cross the focal plane of the platelet aggregates at about 650 s after laser-induced damage. The volume consisted of 1000 scans; an axial z-stack in steps of 1 µm across 100 µm with an average of 10 frames per step. Oversampling and averaging was performed solely to improve the signal-to-noise ratio.

### Blinded test

To test the effect of transient hyperglycemia in an unbiased manner we used randomized blinded i.p. injection of *L*-glucose or *D*-glucose. Prior to each imaging session mice were fasted overnight (18 hr), then injected with *L*-glucose; 2 gm/kg mouse body weight, 20 % (w/v) in saline. Mice injected with *L*-glucose were then imaged and platelet aggregate formation was induced and monitored. An identical second imaging/laser-induced damage session was then conducted with the same mouse, injected with a second unmarked tube which contain either *L*- or *D*-glucose; 2 gm/kg mouse body weight of 20 % (w/v) in saline. Importantly, each imaging session was carried out in a separate venule that branched from the central sinus (**Figure 1A**).

## QUANTIFICATION AND STATISTICAL ANALYSIS

All time-series analysis was done using MATLAB (The Math Works). ROIs were drawn around the laser damage site and the average intensities of the ROI in multiple fluorescence channels were used to construct time series of platelet aggregation, neutrophil aggregation, and serum extravasation. Time series were normalized to the average of the 141 s of data acquired prior to the laser-induced vascular disruption, i.e., these data define F_0_ at each pixel. We then calculate ΔF/F_0_ = F(t)/F_0_ – 1. Volumetric analysis was done on image stacks using Imaris (Oxford Instruments).

To determine the laser power needed to induce thrombosis, a piecewise linear model was fit to the platelet aggregates volume and platelet aggregates fraction as a function of laser energy using the Shape Language Modeling toolbox in MATLAB (D’Errico, 2021). The model had one free interior knot, i.e., the threshold power, and was constrained to be a constant value between 0 and 0.2 µJ of the laser pulse energy.

The two-sided unpaired Student’s t-test was used to compare platelet and neutrophil volumes and volume fractions at 650 s across different experimental conditions.

The one-sided unpaired Wilcoxon rank sum test was used to compare time series of changes in blood component different experimental conditions. For blood serum, we used the average fluorescence of the last 100 s of the relevant times series. For platelet and neutrophil aggregation, the average fluorescence of the first 100 s of the times series after laser damage was compared across different experimental conditions.

A one-way ANOVA was used to perform a meta-analysis of whether platelet aggregates volume and platelet aggregates fraction at 650 s differed significantly across different control and *D-*glucose groups.

## Data sharing

The datasets supporting the current study are available as public links:

Figure 1A: https://www.dropbox.com/s/ldm0phvclda1typ/Fig_1A_JEM.zip?dl=0

Figure 1B-H: https://www.dropbox.com/s/p33e3n3mh2ay8zr/Fig_1B_H_JEM.zip?dl=0

Figure 2: https://www.dropbox.com/s/jvpw2iexc5g3hwq/Fig_2_JEM.zip?dl=0

Figure 3: https://www.dropbox.com/s/bauy44zn9kmhrll/Fig_3_JEM.zip?dl=0

Figure 4: https://www.dropbox.com/s/43912mnlxgtdar3/Fig_4_JEM.zip?dl=0

Figure 5: https://www.dropbox.com/s/sapz75gynxg3ses/Fig_5_JEM.zip?dl=0

Figure 6: https://www.dropbox.com/s/sug0fsnlxz0h9ly/Fig_6_JEM.zip?dl=0

Figure 7: https://www.dropbox.com/s/53iqe6qi6w2ndcm/Fig_7_JEM.zip?dl=0

Figure 8: https://www.dropbox.com/s/bn5a0tuaqlrhbsg/Fig_8_JEM.zip?dl=0

